# Analyzing Single Molecule Localization Microscopy Data Using Cubic Splines

**DOI:** 10.1101/083402

**Authors:** Hazen P. Babcock, Xiaowei Zhuang

## Abstract

The resolution of super-resolution microscopy based on single molecule localization is in part determined by the accuracy of the localization algorithm. In most published approaches to date this localization is done by fitting an analytical function that approximates the point spread function (PSF) of the microscope. However, particularly for localization in 3D, analytical functions such as a Gaussian, which are computationally inexpensive, may not accurately capture the PSF shape leading to reduced fitting accuracy. On the other hand, analytical functions that can accurately capture the PSF shape, such as those based on pupil functions, can be computationally expensive. Here we investigate the use of cubic splines as an alternative fitting approach. We demonstrate that cubic splines can capture the shape of any PSF with high accuracy and that they can be used for fitting the PSF with only a 2-3x increase in computation time as compared to Gaussian fitting. We provide an open-source software package that measures the PSF of any microscope and uses the measured PSF to perform 3D single molecule localization microscopy analysis with reasonable accuracy and speed.

## Introduction

The accuracy of the fitting algorithm is an important factor in the final resolution of single molecule localization based super-resolution microscopy. This accuracy is in turn determined in part by how well the fitting algorithm models the PSF of the microscope. This is particularly important for 3D super-resolution imaging, where the PSF shapes are often not a simple Gaussian function [1-4]. Approaches to accurately modeling the PSF for the purpose of localizing single molecules include PSF correlation [5-7] and pupil function fitting [8]. In the PSF correlation approach, the PSF of the microscope is measured by scanning the objective (or the sample) in Z to capture the shape of the PSF at multiple focal planes. The resulting z stack of PSFs can then be cross-correlated with the image of a single molecule to determine its position. This is computationally expensive as it requires multiple 2D Fourier transforms, followed by further fitting if sub-pixel position information is desired. Alternatively, the molecule’s position can be determined directly by providing an initial estimate for the molecule’s position and computing the fitting error between the molecule’s image and the PSF, then appropriately updating the position estimate. This can be done using a simplex optimization algorithm such as Nelder-Mead [9]. However this class of algorithms, which are used when the derivatives of the error function are not known or are hard to calculate, converge slowly on the optimal solution. An additional complication of this latter approach is the need to interpolate between the measured PSF values in x, y and z.

These problems are largely avoided with a pupil function fitting approach [8, 10, 11], as interpolation is not necessary and the derivatives of the error function can be calculated, enabling the use of more rapidly converging algorithms such as those based on the Newton-Raphson algorithm. In the pupil function approach, the PSF is again measured by scanning the sample containing point emitters (such as beads) in Z, passing through the focal plane of the objective. The measured PSF is then fit to determine the pupil function of the microscope. As the pupil function captures the PSF of the microscope with high accuracy, fitting the images of single emitters to the pupil function allows the positions of these emitters to be determined with high accuracy. However calculating the PSF and its derivatives with respect to the fitting parameters at a particular x, y and z location from the pupil function is computationally expensive, and could be slow unless GPU acceleration is used [8].

Another approach to accurately model the PSF for the purpose of localizing single molecules is to use splines. The most commonly used splines approximate the function of interest using one or more polynomials. This class of splines can be calculated rapidly, and it is also easy to compute their derivatives. 2D cubic splines were used in the DAOPHOT astronomy package to provide higher order corrections to a 2D Gaussian for the purpose of fitting the locations and magnitudes of stars [12], or single molecules [13], the latter using the DAOSTORM algorithm, the adaptation of DAOPHOT for single molecule localization microscopy. B-splines, of which cubic splines are a sub-set, were used previously in localization microscopy to estimate the maximal localization accuracy in x, y and z of an arbitrarily shaped 3D point spread function (PSF) [14]. B-splines were also used in a general method for 3D localization fitting similar to what is presented here [15].

This current work builds on previous work by further exploring the use of splines for rapid and accurate analysis of 3D super-resolution microscopy data based on single molecule localization. In particular, we provide an open-source maximum likelihood estimation (MLE) based cubic spline analysis package, side-by-side comparisons of its performance with elliptical Gaussian fitting, and a demonstration of its broad application by fitting simulated 3D super-resolution microscopy data derived from several different PSFs.

## Materials and Methods

A cubic spline approximates a one dimensional function using piecewise 3^rd^ order polynomials of the form:

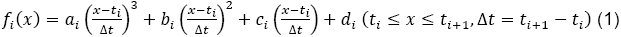

where each *f_i_* is valid on small equal length intervals [*t_i_*, *t*_*i*+1_]. Any function with a continuous first derivative can be approximated with arbitrary accuracy by choosing an appropriate interval size. Here we follow DAOPHOT / DAOSTORM and choose an interval size that is ½ of the image pixel size as a good compromise between accurately modeling the PSF and minimizing the total number of intervals and coefficients.

In Fig 1 we provide a simple 1D example of the cubic spline construction process that we use in this work. We start with a hypothetical measurement of the PSF (gray bars in Fig 1). Next the measured PSF is up-sampled by 2x using 3^rd^ order spline interpolation (red points in Fig 1). Finally the coefficients *a_i_*, *b_i_*, *c_i_*, *d_i_* of the spline (blue line in Fig 1) for each interval are calculated with the constraint that the polynomials *f_i_*(*x*) and their first derivatives *df_i_*(*x*)/*dx* are continuous at the red points, the interval boundaries. We chose to use the additional constraint that the second derivative of the spline is zero at the start of the first interval and at the end of the last interval. We use the approach described in [16] to solve for the coefficients of the (natural) cubic spline.

**Figure 1.**
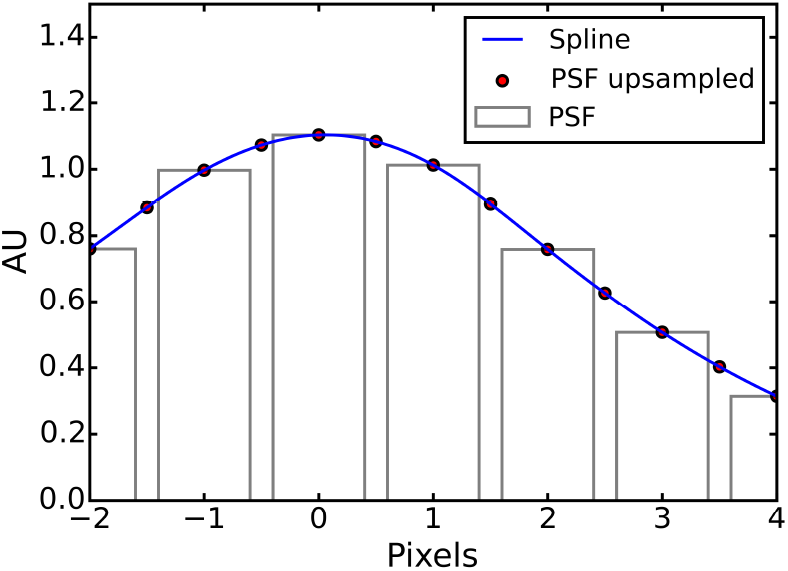
1D Spline example. Each cubic spline interval (the blue line) is determined such that itself and its first derivative are continuous across the red points which are the interval boundaries. The red points are calculated by up-sampling the measured PSF (grey bars) by a factor of two using 3^rd^ order spline interpolation.

Next, we used the following procedure to determine the PSF in three dimensions, first by simulations and then by experiments. To determine the 3D PSF by simulation, we first place a number of emitters on a xy grid with an additional single pixel uniform random offset and generate z scan image stacks of these emitters. Then, using the known emitter locations, images of isolated emitters were cropped out of the image stacks and up-sampled by 2x in xy using the ndimage.interpolation.zoom() function from the Scipy Python package [17] with 3^rd^ order spline interpolation. Next, the up-sampled images of these emitters were aligned to their centroid positions using the ndimage.interpolation.shift() function and averaged to obtain a single image for each z position. Finally, images covering 50 nm z ranges were further averaged together to generate the PSF for the z position that is at the center of the 50 nm range and the average PSFs determined from a series 50 nm intervals were used to create a 3D array that represents the 3D PSF. To measure the 3D PSF in real experiments, small fluorescent beads can be attached to a coverslip. Then z scan movies can be taken with a piezo positioner. In this situation the true emitter locations are not known, but they can be estimated with sufficient precision for this purpose by using another fitting algorithm such as 3D-DAOSTORM [18] because the fluorescent beads typical emit a very large number of photons.

The extension of the cubic spline construction to 3 dimensions is described below. In three dimensions, the cubic spline is:

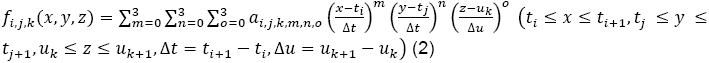

where each *f*_*i,j,k*_ is valid on ½ pixel length intervals [*t_i_*, *t*_*i*+1_] in x, [*t_j_*, *t*_*j*+1_]*in* y and [*u_k_*, *u*_*k*+1_] intervals in z. We start with a 3D PSF that is determined by either simulation or experiments, and then up-sampled such that it has an equal number of elements in x,y and z. This is not strictly necessary but having an equal number of elements for each axis simplifies later computations. We then use a series of 1D cubic spline interpolations in x, y and z to further upsample the PSF, by 4 fold in each dimension, to obtain 64 values of the PSF within each element. These 64 values then allow us to uniquely specify the 3D cubic spline *f*_*i,j,k*_(*x*, *y*, *z*) anddetermine it’s 64 coefficients *a*_*i,j,k,m,n,o*_ in the intervals that correspond to each element of the PSF.

We then use this cubic spline approximation of the 3D PSF to fit the image of individual emitters and determine their 3D coordinates. Based on the 3D cubic spline formula it can be seen that the evaluation of *f*_*i,j,k*_(*x*, *y*, *z*) at a single point will require 128 operations, as compared to 10 operations to evaluate the elliptical Gaussian that is often used for 3D PSF fitting:

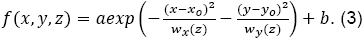

Here we’ve ignored the calculation of the *x*^*m*^*y*^*n*^*z*^*o*^ terms as for fitting the evaluation is always done at the same point in each of the spline intervals. Similarly, we have ignored the calculation of the *w_x_*(*z*) and the *w_y_*(*z*) terms in the Gaussian as these are also the same for each point in the PSF. We wrote a simple C program to time how long it took to perform these calculations, and found that the cubic spline calculation is approximately 2.3x slower. However, we note that this is a best case estimate as the cubic spline calculation is more memory intensive and thus is likely to be somewhat slower in general. However, this rough estimate suggests that the computational expense of the cubic spline fitting will be similar to that of Gaussian fitting.

The highest accuracy estimation of the parameters of the localization of an emitter is typically achieved using non-linear least squares or maximum likelihood estimation (MLE) based fitting. This can be done efficiently if both the error function and its derivatives are easily calculated for any point in the PSF. As described in [19], the error function for MLE fitting is:

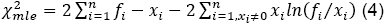

And its derivatives with respect to the fitting parameters (p) are:

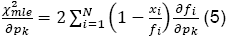

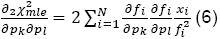

Where we have followed reference [19] and ignored the second derivative terms in the calculation of equation (6).

The cubic spline PSF and its derivatives with respect to the fitting parameters *x_c_*, *y_c_*, *z_c_*, (the 3D coordinates of the emitter), *h* (the height of the PSF), and *b* (the baseline of the PSF) are easy to calculate, and they are:

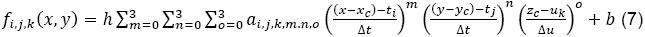

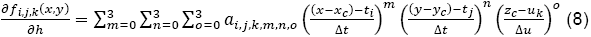

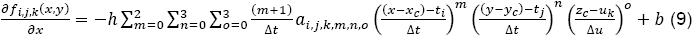

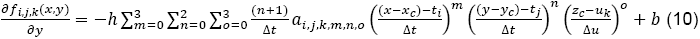

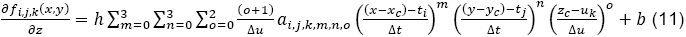

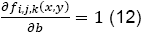

We minimize 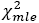 using the above equations and the approach described in reference [19], which is a modified version of the Levenberg-Marquadt algorithm [20]. Given a reasonable starting point the optimal values for *x_c_*, *y_c_*, *z_c_*, *h*, and *b* can be found with high accuracy in just a few (~10) iterative updates for isolated emitters.

In all other respects the approach is the same as that used by 3D-DAOSTORM [18]. The algorithm executes the following pseudo-code on each frame of single-molecule imaging movie until either no new peaks are found, or the maximum number of iterations is reached.

### Algorithm

~~~
1: Load the current image.
2: Perform an initial estimate of the image background.
3: **while not** all converged **and** repeats < max repeats **do**
4:   Subtract current background estimate from the image.
5:   Find localizations in the background subtracted image and add these to the list of all localizations.
6:   Subtract localizations from the image.
7:   **while** iterations < max iterations **and not** all converged **do**
8:        **for each** localization **do**
9:              **if not** converged **then**
**10:                 Add localization to image**
11:                       Levenberg-Marquadt update of localization fitting parameters
12:                       Subtract updated localization from image
13:                       **if** converged **then**
14:                                mark localization as converged
15:  **for each** localization **do**
16:       **if** distance to a brighter neighbor < minimum distance **then**
17:             discard localization
18:             mark all neighbors as unconverged
19:  Subtract localizations from the image.
20:  Estimate background of the localization subtracted image.
21: Save fit parameters of all the localizations.
~~~

Note that this algorithm ignores cross-terms for overlapping localizations. Each localization’s fitting parameters are updated in a context where all the other localizations are fixed. This increases the simplicity of the algorithm implementation at the price of a reduced convergence speed for overlapping localizations.

## Results and Discussion

To test how well a cubic spline with ½ pixel interval size in x and y can capture the shape of non-trivial PSFs we used the pupil function approach [10, 11] to generate 3D PSFs for a purely astigmatic PSF (*Z*_22_ = 1.3) and for the saddle-point PSF described in [3], which have been previously used for 3D super-resolution imaging [1, 3]. For each condition we generated noise free images of the PSF at z positions ranging from -1 um to 1 um in 10 nm steps. Then we determined the coefficients of the cubic spline for this PSF as described in the Materials and Methods section. We found that the maximum pixel wise amplitude difference between the astigmatic PSF and the cubic spline was 2.0% (at z = 0 nm) over the entire 2 um z range (Fig 2). Similarly, we found for the saddle-point PSF that the maximum pixel wise difference was 3.7% (at z = 40 nm). We also tested how well an elliptical Gaussian was able to capture the shape of a purely astigmatic PSF, as elliptical Gaussian is the analytical function typically used in astigmatism-based 3D super-resolution imaging. We calculated the best fit elliptical Gaussian to the astigmatic PSF for each z position in the same 2 um z range and found that the error was somewhat larger than the cubic spline fitting, with a maximum pixel wise difference of 7.7% (at z = 60 nm)

**Figure 2.**
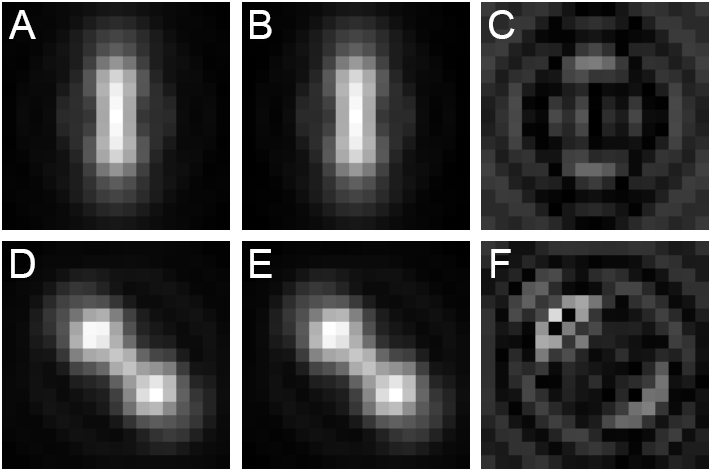
Analytical PSFs and their spline representations for the astigmatic PSF and saddle-point PSF. (A) Astigmatic PSF at z = -420 nm. (B) Cubic spline representation of (A).(C) Difference between (A) and (B) multiplied by 100. (D) Saddle-point PSF at z = -420 nm. (E) Cubic spline representation of (D). (F) Difference between (D) and (E) multiplied by 100.

Next we investigated whether the fact that the cubic spline more accurately captured the shape of a purely astigmatic PSF resulted in improved fitting performance. First we created a calibration data set by placing many emitters on a xy grid with an additional single pixel uniform random offset and then scanning the z position of the emitters with a 10 nm interval. We then used these calibration movies to determine the coefficients of the cubic spline representation of the PSF and also to determine the *w_x_*(*z*) and *w_y_*(*z*) calibration curve for elliptical Gaussian z fitting. Then a simulated movie was created with 3000 photons per localization and a constant background of 100 photons. In the test movie, we chose the z position of the emitters to range from -300 nm to 300 nm, as it has been noted previously that the localization precision of the astigmatism approach depends on the z position of the emitter and that relatively high localization precisions have been achieved experimentally in the z range of -300 to 300 nm [1]. These movies were created using a Poisson noise model, with no camera read noise, 600 nm emission wavelength and a 160 nm pixel size. The test movie was analyzed by fitting the single-emitter images to the cubic spline representation and elliptical Gaussian representation of the astigmatic PSF (Fig 3). The fitting error in x, y and z for each z value was determined as the average distance in x, y and z between the localizations determined by the fitting and their nearest ground-truth emitter positions.

**Figure 3.**
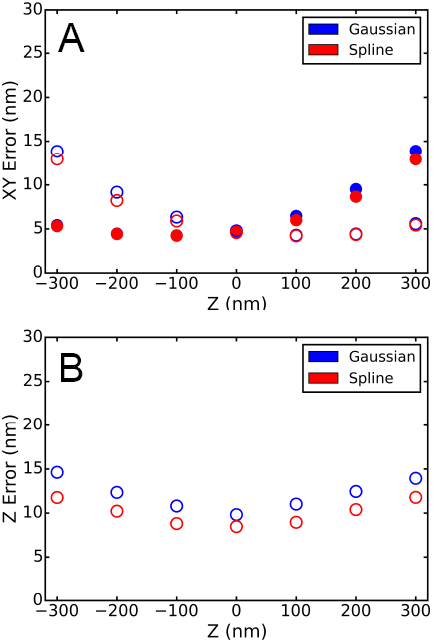
Astigmatic PSF fitting accuracy. (A) Average xy fitting error as a function of z for cubic spline versus elliptical Gaussian fitting. Ellipitical Gaussian fitting were performed with the 3D DAOSTORM algorithm that we reported previously [18]. The filled points are the error in x and the hollow points are the error in y. Each data point is the average of multiple independent z positions in the 100 nm range centered on the data point. (B) Average z fitting error as a function of z.

The x and y localization error was pretty similar for both approaches, with cubic spline fitting giving 5-10% better accuracy at the largest z offsets. The z localization accuracy of cubic spline fitting however was noticeable better, with an average improvement of about 20% at larger z offsets. We note that in real experiments, the PSF often deviates from the ideal PSF derived from the pupil function due to non-ideal optics and optical alignment errors, which can lead to systematic errors in the determination of the x, y and z positions of the emitters when a simple elliptical Gaussian function is used to fit the PSF [21, 22]. Since the cubic spline representation can capture these experimental PSFs with arbitrarily high accuracy given a sufficiently small interval size, we expect cubic spline fitting to provide a greater improvement in the localization accuracy in experiments.

The average amount of time that it takes to analyze a single image with the cubic spline approach as a function of emitter density is shown in Fig 4. Timing information for the elliptical Gaussian fitting (using the 3D-DAOSTORM algorithm) is provided for comparison. All timing measurements were performed on a laptop computer with an Intel Core i7 CPU running at 2 GHz. Simulated movies (41 um × 41 um field of view) were generated with emitters randomly located in x and y in a +−500 nm z range at densities of 0.03, 0.1 and 0.3 per um^2^ (41, 136 and 408 emitters respectively). The PSF that we used in these simulations was an elliptical Gaussian whose width in x and y varied as a function of z. To reduce the variance in the timing results, we increased the number of frames of the movies until they took at least 4 seconds to analyze regardless of the emitter density. As 3D-DAOSTORM and the cubic spline algorithm share most of the other components of their analysis pipeline, the timing differences reported are primarily due to differences in how long the localization fitting step takes. At lower emitter densities the overall analysis time is more dependent on how long the various non-fitting steps, such as reading the image from disk and background estimation, take, therefore the computation time of the two algorithms is similar. At higher emitter densities the fitting step takes a larger fraction of the total time, and hence the cubic spline approach takes a proportionally longer time. These results are also consistent with our estimates for the relative computation cost of the two approaches described earlier.

**Figure 4.**
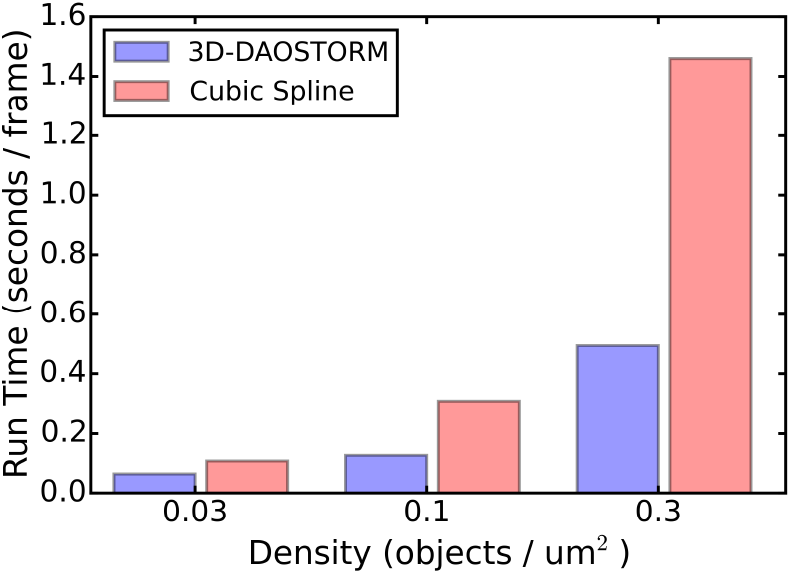
Fitting speed comparison. A comparison of the time it takes to analyze a single frame (256 × 256 pixels) of a simulated movie with indicated emitter densities using 3D-DAOSTORM or the cubic spline approach.

It is worth noting that the fitting time will depend on the number of pixels that the PSF covers. In the above test, we used a spline that covers 14 × 14 pixels in x and y. This is a reasonable size for most PSFs, but a larger size may be necessary for PSFs with a larger spatial extent or for microscope setups with a smaller pixel size. We found a size of between 12 and 16 pixels (in each dimension) was sufficiently large to work well for the PSFs that are typically used in single-molecule imaging and our pixel size of 160nm.

An important advantage of the cubic spline approach is that it can be used to fit arbitrarily shaped PSFs, which makes it widely applicable to many different 3D localization methods using various engineered PSF shapes. Here, we demonstrate the versatility of the cubic spline approach by using it to analyze simulated data from two additional types of PSFs that have been used for super-resolution imaging, the double helix PSF [2] and the saddle-point PSF [3]. We generated two kinds of simulated movies using these PSFs, a high photon count movie for measuring the PSF and determining the spline and a low photon count movie similar to typical single-molecule localization conditions. The saddle-point PSFs were generated using the pupil function approach [10, 11], and we rotated the saddle-point PSF by 45 degrees relative to the orientation reported in [3] so that the PSF stretches along the x and y axes. The double helix PSF was obtained from the 2016 SMLM localization challenge web-site [23] and spline interpolation was used to position it at arbitrary x, y and z values. In the movies simulating single molecules, we used 3000 photons per localization, a constant background of 100 photons, and a camera pixel size of 160 nm. The x, y and z errors were again determined as the average distance in x, y and z between the localizations determined by the fitting and their nearest ground-truth emitter positions and shown in Figure 5. From these error plots, it is evident that the cubic spline approach can fit any of these PSFs with high accuracy.

**Figure 5.**
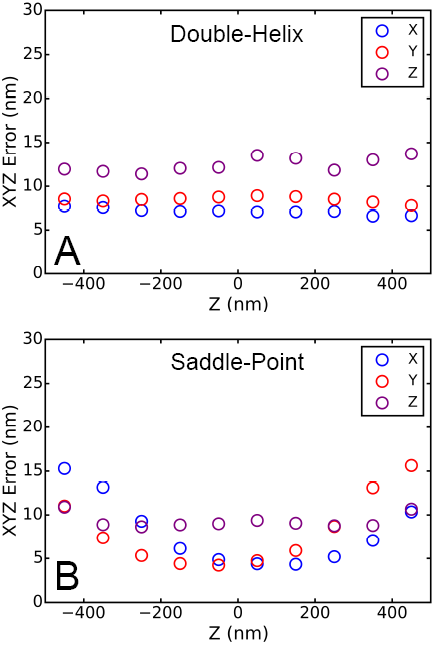
PSF localization accuracy. Localization errors in x, y and z are plotted as a function of z. (A) double helix PSF (B) saddle-point PSF.

We note that the localization errors reported in this work are all determined by fitting the relatively ideal, simulated images of single molecules with the same photon numbers for each PSF. In real experiments, the photon loss associated with generating relatively complex PSFs, for example by using spatial light modulators or deformable mirrors, also needs to be taken into account when estimating the final localization error.

## Conclusions

In this work, we demonstrate that a cubic spline approach provides a 5-10% improvement in x and y localization accuracy and up to a 20% improvement in z localization accuracy relative to elliptical Gaussian fitting for a purely astigmatic PSF. The cubic spline approach has the additional advantage of being very flexible, as it is easily adapted to fit any experimental realizable PSF. It is also a practical choice as it is only ~2-3x slower than a Gaussian fitting approach at typical emitter densities encountered in super-resolution imaging experiments. To make it easier for others to explore this approach to single-molecule localization analysis, our implementation of this algorithm Docker images, instructions and an example of its use are available on Github [24].

## Acknowledgements

We thank YoavShectman for providing the pupil function for the saddle-point PSF. This work was supported in part by the National Institute of Mental Health Conte Center Program (P50MH094271). X.Z. is a HHMI investigator.

## References

1. Huang B, Wang W, Bates M, Zhuang X. Three-Dimensional Super-Resolution Imaging by Stochastic Optical Reconstruction Microscopy. Science. 2008;319(5864):810–3.

2. Pavani SRP, Thompson MA, Biteen JS, Lord SJ, Liu N, Twieg RJ, et al. Three-dimensional, single-molecule fluorescence imaging beyond the diffraction limit by using a double-helix point spread function. Proc Natl Acad Sci U S A. 2009;106(9):2995–9.

3. Shechtman Y, Steffen JS, Backer AS, Moerner WE. Optimal Point Spread Function Design for 3D Imaging. Physical Review Letters. 2014;113(133902):1–5.

4. Jia S, Vaughan JC, Zhuang X. Isotropic three-dimensional super-resolution imaging with a self-bending point spread function. Nature Photonics. 2014;8:302–6.

5. Juette MF, Gould TJ, Lessard MD, Mlodzianoski MJ, Nagpure BS, Bennett BT, et al. Three-dimensional sub-100 nm resolution fluorescence microscopy of thick samples. Nature Methods. 2008;5:527–9.

6. York AG, Ghitani A, Vaziri A, Davidson MW, Shroff H. Confined activation and subdiffraction localization enables whole-cell PALM with genetically expressed probes. Nature Methods. 2011;8(4):327–33.

7. Mlodzianoski MJ, Juette MF, Beane GL, Bewersdorf J. Experimental characterization of 3D localization techniques for particle-tracking and super-resolution microscopy. Optics Express. 2009;17(10):8264–77.

8. Liu S, Kromann EB, Krueger WD, Bewersdorf J, Lidke KA. Three dimensional single molecule localization using a phase retrieved pupil function. Optics Express. 2013;21(24):29462–87.

9. Nelder JA, Mead R. A simplex method for function minimization. The Computer Journal. 1965;7(4):308–13.

10. Hanser BM, Gustafsson MGL, Agard DA, Sedat JW. Phase retrieval for high-numerical-aperture optical systems. Optics Letters. 2003;28(10):801–3.

11. Hanser BM, Gustafsson MGL, Agard DA, Sedat JW. Phase-retrieved pupil functions in wide-field fluorescence microscopy. Journal of Microscopy. 2004;216:32–48.

12. Stetson PB. DAOPHOT: A Computer Program for Crowded-Field Stellar Photometry. Publications of the Astronomical Society of the Pacific. 1987;99:191–222.

13. Holden SJ, Uphoff S, Kapanidis AN. DAOSTORM: an algorithm for high-density super-resolution microscopy. Nature Methods. 2011;8(4):279–80.

14. Tahmasbi A, Ward ES, Ober RJ. Determination of localization accuracy based on experimentally acquired image sets: applications to single molecule microscopy. Optics Express. 2015;23(6):7630–52.

15. Kirshner H, Vonesch C, Unser M. Can localization microscopy benefit from approximation theory? 2013 IEEE 10th Internation Symposium on Biomedical Imaging. 2013:588–91.

16. Anton H, Rorres C. Elementary linear algebra with applications: Wiley; 1987.

17. Jones E, Oliphant T, Peterson P. SciPy: Open source scientific tool for Python 2001 [cited 2016 2016-06-06]. Available from: www.scipy.org.

18. Babcock HP, Sigal YM, Zhuang X. A high-density 3D localization algorithm for stochastic optical reconstruction microscopy. Optical Nanoscopy. 2012;1.

19. Laurence TA, Chromy BA. Efficient maximum likelihood estimator fitting of histograms. Nature Methods. 2010;7(5):338–9.

20. Marquardt DW. An algorithm for Least-Squares Estimation of Nonlinear Parameters. Journal of the Society for Industrial and Applied Mathematics. 1963;11(2):431–41.

21. Proppert S, Wolter S, Holm T, Klein T, van de Linde S, Sauer M. Cubic B-spline calibration for 3D super-resolution measurements using astigmatic imaging. Optics Express. 2014;22(9):10304–16.

22. Carlini L, Holden SJ, Douglass KM, Manley S. Correction of a Depth-Dependent Lateral Distortion in 3D Super-Resolution Imaging. PLOS One. 2015;e0142949:1–15.

23. SMLM 2016 [cited 2016]. Available from: http://bigwww.epfl.ch/smlm/#&panel1-2.

24. Storm-Analysis [cited 2016]. Available from: https://github.com/ZhuangLab/storm-analysis.

